# Expression Microdissection for use in qPCR based analysis of miRNA in a single cell type

**DOI:** 10.1101/2022.06.24.497524

**Authors:** Ana E. Jenike, Brady Bunkelman, Kira A. Perzel Mandell, Cliff Oduor, Deborah Chin, Devin Mair, Katharine M. Jenike, Deok-Ho Kim, Jeffrey A. Bailey, Miriam H. Rafailovich, Avi Z. Rosenberg, Marc K. Halushka

## Abstract

Cell-specific microRNA (miRNA) expression estimates are important in characterizing the localization of miRNA signaling within tissues. Much of this data is obtained from cultured cells, a process known to significantly alter miRNA expression levels. Thus, our knowledge of *in vivo* cell miRNA expression estimates is poor. We previously demonstrated expression microdissection-miRNA-sequencing (xMD-miRNA-seq) as a means to acquire *in vivo* estimates, directly from formalin fixed tissues, albeit with limited yield. Here we optimized each step of the xMD process including tissue retrieval, tissue transfer, film preparation, and RNA isolation to increase RNA yields and ultimately show strong enrichment for *in vivo* miRNA expression by qPCR array. These method improvements, including the development of a non-crosslinked ethylene vinyl acetate (EVA) membrane, resulted in a 23-45 fold increase in miRNA yield, depending on cell type. By qPCR, miR-200a was increased 14-fold in xMD-derived small intestine epithelial cells, with a concurrent 336-fold reduction in miR-143, relative to the matched non-dissected duodenal tissue. xMD is now an optimized method to obtain robust *in vivo* miRNA expression estimates from cells.

## Background

microRNAs (miRNAs), small noncoding RNAs, are essential regulators of mRNA translation. miRNAs are intrinsic regulators of cellular physiology and have been linked to multiple pathologies through expression level changes in tissues^1-3^. However, tissues are diverse landscapes of multiple cell types all contributing to the miRNAome of that tissue^4-7^(New Arun paper). The small intestine, for instance, has numerous discrete cell types of epithelial, endothelial, and inflammatory lineages^8^. It would be ideal to independently identify the miRNA expression pattern of each cell type. Cell culture is frequently used for this task, however, culturing and passaging cells dramatically alters miRNA expression patterns^9^. Therefore, the miRNA expression, when obtainable from cell culture, is not a perfect proxy to *in vivo* cellular expression patterns. Additionally, many cell types such as cardiomyocytes and neurons do not grow effectively in culture. There is currently a need for an effective and efficient method to isolate cells directly from tissues to best approximate *in vivo* miRNA expression patterns.

Two commonly used methods, laser capture microdissection and flow cytometry, exist for quantifying expression levels of miRNAs within distinct cell subsets. Laser capture microdissection is perhaps the most global and accurate, but it is best performed with frozen tissues which are limiting for human studies^10^. Also, it is laborious and requires technical expertise and expensive machinery. Flow cytometry with cell capture, like laser capture, requires expensive equipment and is somewhat limited by the available antibodies to mark different non-immune cell types^11^. The method to dissociate cells has long preparation times during which cell stress increases expression alterations^5^. These concerns highlight the necessity of new microdissection techniques that allow for the retrieval of the near *in vivo* miRNA expression signatures.

Expression microdissection (xMD) is a method to rapidly and cost-effectively microdissect cells directly from tissue on glass slides^12,13^. We previously introduced xMD-miRNA-seq as an extension of the method to specifically obtain the miRNA signature of intestinal epithelial cells^14^. In our initial development of xMD-miRNA-seq, we noted an 80-fold reduction in RNA from an unprocessed slide to the final xMD membrane-obtained material during the xMD steps as well as a low percentage of miRNA reads from the sequencing library preparation. Here we present an optimized method of xMD to assay miRNAs where we have substantially optimized the RNA collection for the purpose of qPCR array. We demonstrate a step-by-step approach to evaluate each facet of the method to increase specificity and RNA yield. These improvements have increased the opportunities to utilize xMD on both common and rare varieties of cell types to best understand the *in vivo* expression patterns of miRNAs in health and disease.

## Methods

### Procurement of Human Tissue

Sections of duodenum (small intestine) were procured from pancreatoduodenectomy specimens and heart tissue was collected from an orthotopic heart transplant case in an expedited fashion in the surgical pathology suite at The Johns Hopkins Hospital. Johns Hopkins Investigational Review Board (IRB) approval was given and patient consent was obtained for use of these tissues. Experiments were performed in accordance with our guidelines including anonymization upon receipt. Specimens were formalin fixed for 24 hours, followed by standard processing and paraffin embedding. Four micron (μm) sections were placed on Superfrost Plus slides (Fisherbrand, Cat No. 12-550-15) and stored at-80°C until use.

### Evaluating RNA loss at different steps in the IHC protocol

Prior to optimizing the full protocol, we performed a general analysis of multiple steps of the original IHC protocol to determine key steps that caused significant RNA loss. Slides underwent one or more steps of baking, antigen retrieval by either high temperature antigen retrieval (HTAR) or proteinase K (15 minutes [min]), primary antibody staining (AE1/AE3, Bio SB, Cat No: BSB 5432), and or Poly linker (secondary antibody) staining. Following certain steps, heart tissue slides were scraped using a razor blade and tissue was collected for RNA processing as described below. Variations including either 5 or 15 min of HTAR, either EDTA or citrate for the HTAR, and the presence or absence of one of two RNAse inhibitors (1 – Millipore Sigma, Protector RNAse Inhibitor, Cat. No. 3335399001; 2 – NEB, RNAse Inhibitor, M0314S). The RNA was then extracted from the tissue using the miRNeasy kit (Qiagen Cat. No. 217084). The amount of RNA was evaluated using qPCR, for hsa-miR-133 relative to a Cel-miR-39 spike-in.

### Global Expression Microdissection Protocol

The complete, final version of the protocol is given herein, with the experimentally modified steps noted as (**A-G**). The slides were deparaffinized before staining by heating at 60°C for 20 min (Thermobrite StatSpin system) and then washed in 3 xylene baths for 5 min each (Macron, ACS grade), 2 ethanol baths for 3 min each (Pharmco, Cat No: 111000200), followed by 3 min in 90%, and 3 min in 80% ethanol respectively. The slides were placed in a citrate solution (Bio SB, Cat No: BSB 0020) and antigen retrieved using a high pressure high temperature (HTAR) method with a pressure cooker (Cuisinart). The entire HTAR process included 20 min of ramp up time, 1 min at full pressure and temperature and 7 min of cool down time (**A**). The slides were treated with peroxide blocker (Bio SB, PolyDetector Plus) for 5 min. For epithelial cell staining, the primary antibody was anti-AE1/AE3 (anti-pancytokeratin) (Bio SB, Cat No: BSB 5432) at a 1:100 dilution for 45 min. For endothelial cells, anti-CD31 (Bio SB, Cat No: BSB 5223) was used at a 1:75 dilution for 60 min. To each antibody solution, 15μl of RNASecure (Thermo Fisher, Cat No. AM7006) per ml was added (**B)**. Primary antibodies were washed off with immuno-wash and treated with Poly linker and Poly HRP for 15 min each with washes in between. The slides were treated with the chromogen 3,3’-Diaminobenzidine (DAB) for 10 min (Biocare Medical, Cat No: DB801) (**C**). The slides were washed again with immuno-wash (Bio SB PolyDetector Plus) then dehydrated with serial ethanol baths of increasing concentrations followed by 3 xylene baths. No counterstaining or coverslipping was performed. After staining, slides were stored at -80°C until use.

The xMD nucleic acid material isolations from tissues were performed using a SensEpil flash lamp (HomeSkinovations, AS101500A), a food saver storage system (FoodSaver, v3835), and fullerene impregnated Ethylene Vinyl Acetate (EVA) (**D**). Stained slides were covered with an initial trimmed EVA membrane placed on the tissue and pressed down using a wooden dowel. A second EVA membrane was then sealed against the slide using the FoodSaver vacuum system to tightly oppose the two (**E**). Then a wetted western blot sponge (Thermo Fisher, Cat No. EI9052) was placed on top of the vacuum bag. The flash lamp was placed on top of the sponge and flashed 5 times at the intensity 4 (of 5) over a white shiny background (**F, G**). The vacuum bag was opened, the slide/EVA removed and the EVA membrane, containing the transferred tissue was gently detached and placed in a 1.5 ml microcentrifuge tube for digestion. After xMD, two membranes were placed in each sample tube and frozen at -80°C overnight, allowing a freeze/thaw to occur which improved the dissolution of the membrane. 300 μl of Protein Kinase Digestion (PKD) buffer (Qiagen, Cat. No. 169021771) was added to sample tubes, to cover the membranes, more was added if the membranes were not covered. Membranes were incubated with 10 μl proteinase K at 56°C for 30 min, followed by 15 min at 80°C to deactivate the enzyme. Afterwards, the samples were treated with DNase for 15 min at room temperature. Then, membranes were incubated for 5 min with one volume (generally 310 μl) of phenol:chloroform (Sigma, Cat. No. P3803). The membrane backers were removed from the tube with tweezers leaving just the mostly dissolved EVA membranes and tissue behind. The samples were incubated at room temperature for 1 hour, under aggressive agitation then centrifuged for 30 min at max speed (16,000 rpm), before removing the layer of chloroform. A 15 min incubation with 20 μl of proteinase K at 56°C, was followed by a 5 min incubation on ice. Another volume of phenol:chloroform was added and the samples were incubated for 5 min at room temperature. The samples were centrifuged for 30 min at max speed (16,000 rpm), in a tabletop centrifuge, and the supernatant (aqueous phase) was transferred to a new tube. Then one volume of isopropyl alcohol (VWR, Cat. No. 0918) was added to the aqueous phase and incubated at -20°C for 3 hours to overnight. The samples were centrifuged for 30 min at 16,000 rpm. The supernatant was discarded and the pellet washed with ethanol twice. The RNA was then resuspended in 20 μl of RNAse free water (Invitrogen, 10977-015). The samples were cleaned with the Zymo Research Clean & Concentrator-5 Kit (Cat no. R1015), including the DNAse step. The RNA was stored at -80°C until use.

### qPCR method for verifying optimization steps

cDNA was synthesized using miScript II RT Kit (Qiagen, Cat No. 218161), according to manufacturer specifications. qPCR was performed using miScript assays (Qiagen, Cat No: 218075) The cDNA was diluted to approximately 40 ng/ul. A master mix was made for all wells with SYBR green master mix with universal primers. Then the mix was aliquoted for each primer, hsa-let-7a, hsa-miR-101 (or hsa-miR-222), hsa-miR-128, hsa-miR-133 and cel-miR-39 as appropriate for the experiment. The specific primers were added to the smaller master mixes. For each treatment, the qPCR was performed in quadruplicate. In each well there was 160 ng cDNA at a concentration of 40 ng/μl. The reaction volume was 25 μl. The thermocycler used the following program: 40 cycles of 95°C for 15 seconds, 55°C for 30 seconds and then 70°C for 30 seconds. The ΔΔ Ct value was calculated relative to the spike in and normalized to the slides that did not undergo IHC.

### qPCR arrays for verifying overall optimization

Human miRNA TLDA cards (Thermo Fisher, Cat No. 44913) were used for global analysis of 384 miRNAs. The RNA extraction was performed as above. The RT was performed using Thermo Fisher Megaplex Primer Pools (Cat no. 4444750) and TaqMan MicroRNA Reverse Transcription Kit (Cat No. 4366596). The RT was followed by a preamplification step using Taqman PreAmp Master Mix (Cat. No. 4488593) and Megaplex PreAmp primers (Cat No. 444750) The array card was then prepared using TaqMan Universal Master Mix II (Cat. No. 444043) and run on a Quantstudio 21k, using the standard protocol (Thermo Fisher, publication 4399813).

Data was analyzed in Microsoft Excel. All controls were checked to assess the quality of work. The data was normalized using the average of three miRNAs known to have steady expression in most cell types (miR-21, miR-22, and miR-103)^15^. All three were assessed to be expressed evenly in all cell types. The CT values for all three miRNAs were averaged for both the tissue scrape and xMD samples. This value was then used to calculate the ΔCT value for all 384 miRNAs in the plate, excluding miRNAs that did not have amplification on any of the sample plates. The 2^-ΔΔ*CT*^ value was then calculated between the tissue scrapes and xMD samples. The fold change was then graphed for each of the miRNAs.

### Protocol modifications

#### A: Varying antigen retrieval time

To assess the role of HTAR in RNA stability, the time at full temperature and pressure was varied. These times were 1 min, 5 min, and 15 min. Each of these HTAR methods had 27 min of heat up/cool down time, thus they represented 28, 32, and 42 min of full experimental time. These experiments were performed with the standard IHC method and evaluated by qPCR as described above (N=4 each arm).

#### B: Evaluating RNAsecure

The RNAse Alert system (Thermo Fisher, Cat no. AM1964) uses a fluorophore-based RNAse sensitive marker to identify RNAse activity. All experiments were performed using 96-well flat-bottomed black plates (Costar, Corning) in a CLARIOstar Monochromator Microplate Reader (BMG Labtech). The CLARIOstar was set to 37°C, a gain of 1400, focal height of 10 mm, excitation/emission wavelengths of 490 nm/520 nm, orbital averaging on at 3 mm, top-optic, 8 flashes per well, and double orbital shaking prior to plate reading at 200 rpm. Reagents related to the IHC steps were tested for intrinsic RNAse activity in either duplicate or triplicate. Four separate antibodies: AE1/AE3 (Diagnostic Bio Systems, Cat. No. Mob 190), Podocin (Sigma Aldrich, P0372-200uL, 096M4797V,), Claudin 8 (R85351 Atlas HPA060605), and CD34 (Sigma Aldrich, HOA036723-100uL, R33262) were evaluated. Additionally, Millipore ddH20, the peroxidase block, poly link, poly horseradish peroxidase (HRP), diaminobenzidine (DAB) (Cardassian DAB, Biocare Medical, Cat. No. DS900) substrate, wash buffers, nuclease free water and controls were evaluated. The RNAse Alert reagent was used according to the manufacturer’s recommendations with measurements taken every 15 min for up to 1 hour.

RNAsecure RNAse Inactivation Reagent (Thermo Fisher) and 0.1% DEPC were evaluated for reducing RNA loss between sets of slides either treated or not treated with the reagent during the primary antibody step of the IHC method. These were compared to slides that only underwent the antigen retrieval step (N=4 each).

#### C: Comparing DAB and DSB chromogens

Comparisons of chromogens that absorbed more light versus a chromogen with toxicity to nucleic acids were made. The IHC was performed as initially described. Slides were stained with either DAB (Biocare Medical, cat No: DBC859l10) or Deep Space Black (DSB, Biocare Medical, BRI4015) (N=4 each).

#### D: Elvax membrane creation and testing

The previously used 3M EVA membranes are insoluble in phenol:chloroform, due to cross-linking, resulting in reduced RNA recovery and isolation. To overcome this, a non-cross-linked membrane was developed using spin coating. ELVAX 410 (Univar Solutions) was coated onto a PETE film (McMaster and Carr, Cat. No. 8567K14) backer using a Lurral spin coater. The backer was prepared by cutting to size, washing in ethanol, then soaking in hexane for 24 hours before drying for 3 hours. The EVA solution for spin coating was made by saturating hexane with fullerene 60 for 3 hours, and then removing the undissolved fullerene. EVA was added at 8% EVA by weight to hexane by volume. The EVA solution was made by heating the solution to 85°C for 5 hours and stirring, until the solution was entirely clear and light purple. The solution was kept at 85°C for the duration of the method.

A two-layer method was used to coat the backer with EVA. A 6 cm diameter backer was treated with 2.5 ml of liquid ELVAX while spinning at 400 RPM. The RPM was then increased to 700 RPM for 30 seconds, 2.5 more ml were added and the new membrane spun for 60 more seconds at 700 RPM. The membrane was then baked at 60°C for 45 min, and allowed to cool for 30 min before being placed in a glass petri dish for storage. To make a membrane without fullerene the same steps were followed but without the addition of fullerene.

The thickness of Elvax 410 membranes was then measured using two methods. The first was a gravimetric method, which was calculated using the EVA density and area of the Elvax 410 membrane. The second was a filmetric method, using a Filmetrics F60-t.

Specificity of the fullerene membranes was tested using human heart slides stained for intercalated disks. The IHC was performed as described above, except with the NCAD antibody (Thermo Fisher, Cat No. MA1-91128) with a dilution of 1:2000. The small yet dispersed nature of intercalated disks in heart tissue allowed for testing visualization of tissue pulled from around the target material. Once the tissue was stained, slides underwent standard xMD, with either fullerene (N=6) or non-fullerene impregnated membranes (N=6). Images of the post dissection membranes were taken and pixels assigned as background, disk or tissue using ImageJ^16^. The percentage of tissue on the membrane that was intercalated disk was calculated.

IHC was performed using AE1/AE3. xMD was performed in the same method as above on either the 3M EVA membrane, the ELVAX 410 EVA membrane, or ELVAX 410 EVA fullerene membrane. The miRNA was then extracted from the membranes with a phenol:chloroform extraction, and the concentrations were compared using a Qubit 4. This was done with five samples for each type of membrane.

#### E: xMD specificity optimization

The process of enhancing xMD specificity was optimized from the initial protocol, with four alterations. All the modifications were evaluated separately, with a final step including all optimizations. The first specificity alteration was the temporary apposition of an EVA membrane to the stained slide, pre-Flash lamp to allow for the removal of loose nucleic acid material from the top of the sample. The second cleaning method was to blow loose nucleic material from the stained slide. High pressure N2 gas, at 95 psi, was blown across each slide with a slow moving nozzle of approximately 0.1 cm diameter for 50 seconds. An additional optimization step was to reduce the amount of flash lamp energy. This was done by decreasing the number of flashes each slide was exposed to for a total of five flashes at the second highest intensity equaling approximately 1724 kJ.

The final optimization method to increase specificity was to perform the xMD on an ice block allowing the samples to cool to close to 0°C. Within a freezer bag, the slides and EVA membrane were cooled for 15 min on an ice block, prior to using the flash lamp. miRNA from all samples was extracted using the standard phenol:chloroform method described above for xMD. qPCR was then performed using the miRcury-LNA kit from Qiagen, following their published methods. The primers used were for miR-143 and miR-192, known to be specific to mesenchymal and epithelial cells respectively. The proportions of the miRNAs were compared across methods and to the standard xMD method.

#### F: xMD background and flash number

The SensEpil flash lamp has 5 levels of flash intensity and each slide could be flashed between 1 and 5 times. Various numbers of flashes and intensities were assessed to compare the capture and specificity of 3M Elvax membrane pulls. Small intestine slides were stained with AE1/AE3 and prepared for microdissection using the standard protcol. These were tested at the highest intensity (5) with 1 flash, 3 flashes and 5 flashes. Then dissections were tested at the lowest intensity (1) with 3 or 5 flashes. Five flashes is approximately 1724 kJ of total energy transfer.

#### G: Background color comparisons

A comparison of five different backgrounds (black, matte white, glossy white, mirror, and no background) were made. Certain differences were qualitatively apparent and others were evaluated using a digital method (HALO) to count the number of pigmented pixels per shared unit area using digital images of EVA membranes with transferred material.

## Results

### Identifying the IHC protocol steps with the largest impact on RNA degradation

There are a number of IHC steps in the xMD process which might be responsible for the loss of RNA through degradation. Therefore, we evaluated these steps by comparing levels of the miRNA let-7a normalized to a cel-miR-39 spike-in, by qPCR, stopping the method after different steps of the IHC process and with varying conditions. Three different antigen retrieval techniques were evaluated on cardiac tissues: proteinase K (PK) digestion, HTAR with EDTA buffer, and HTAR with citrate buffer. The method was stopped at points including just after HTAR, after primary antibody or after the complete IHC protocol. At each stopping point, the slides were scraped of tissue material, RNA was isolated, and qPCR was performed. All miRNA Ct results were compared via fold change to the control unbaked slides. PK digestion for antigen retrieval showed a slight increase (1.4 fold) in let-7a expression (Fig. 1a). Experiments stopping after HTAR with EDTA or citrate showed between a 2- and 15-fold loss of let-7a depending on the length (5 or 15 min) of antigen retrieval (Fig. 1a). A full IHC protocol, using EDTA HTAR resulted in a ∼80 fold loss of let-7a. This was abrogated through the use of two different RNase inhibitors that reduced the loss to ∼40-50 fold (Fig. 1a). Of note, the proteinase K method released more miRNA initially, but this miRNA appeared to be lost in later steps. This experiment indicated that the more significant loss of RNA occurred during the staining steps, rather than the antigen retrieval steps, but that both steps could be optimized to reduce RNase activity.

**Figure 1.**
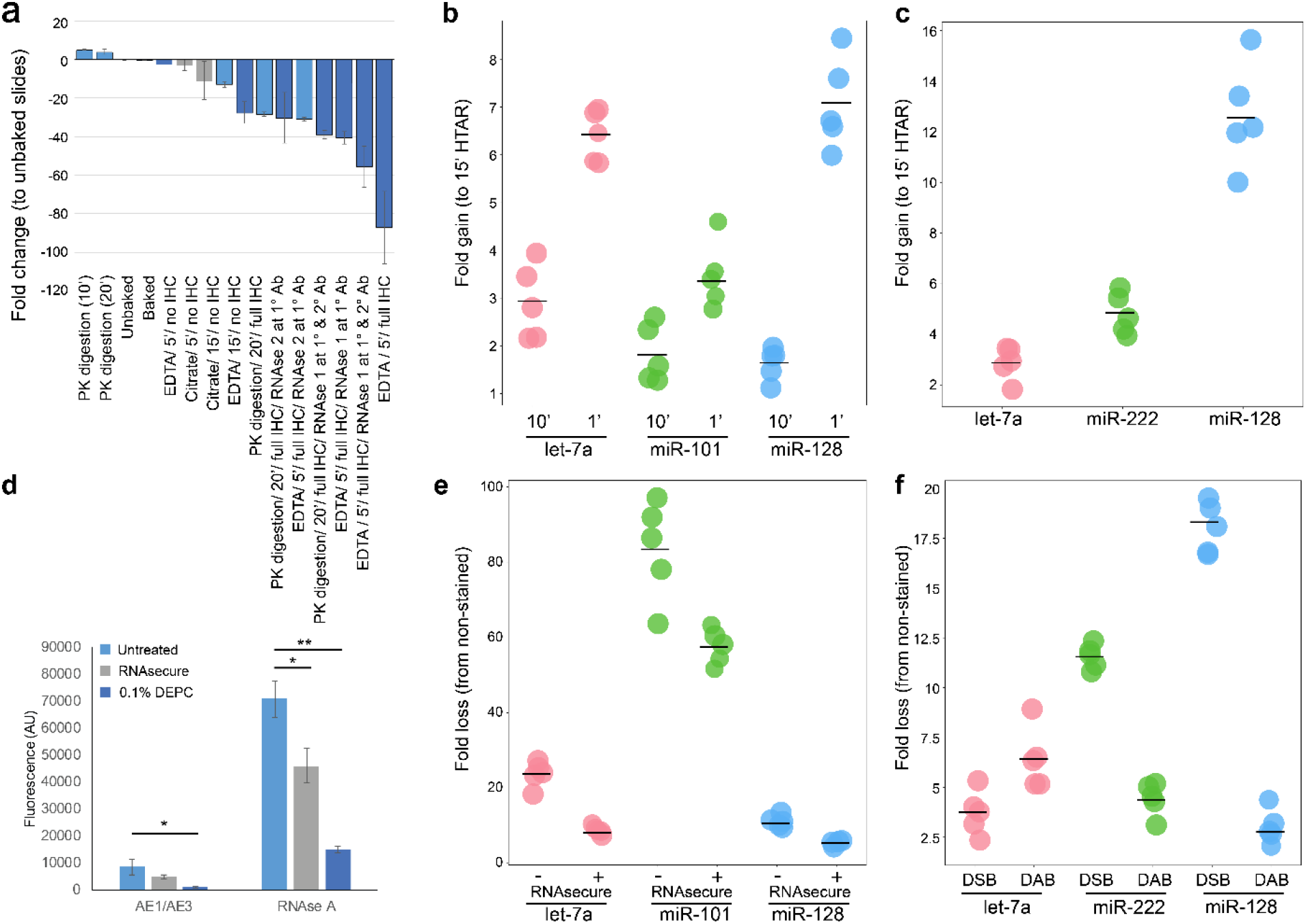
The RNA fold loss of multiple steps was compared.(**a**) Comparison of RNA fold loss between steps of IHC. The fold change is loss of let-7a expression relative to an unexperimented upon tissue scrape from a matched slide. Antigen retrieval methods and the use of RNAse inhibitors (1 – Millipore Sigma / 2 – NEB) were evaluated. >85 fold RNA loss was observed with the standard protocol. (**b**) Increase of miRNA expression in XMD-obtained AE1/AE3+ cells with shorter HTAR lengths (1 & 10 min vs 15 min). (**c**) Increase of miRNA expression in XMD-obtained CD31+ cells with a shorter HTAR length (10 min vs 15 min). (**d**) RNAse A and 0.1% DEPC both reduced the amount of RNAse activity (measured in arbitrary fluorescence units) in AE1/AE3 stained slide material or in material with an RNAse A spike-in. * p<0.01, ** <0.001. (**e**) Whole slides either untreated or treated with RNAsecure during IHC were compared to slides not treated. miRNA loss was greater for three representative miRNAs, Let-7a, miR-101, miR-128 when RNAsecure was not used. (**f**) Slides treated with Deep Space Black (DSB) or DAB were compared to slides not treated with a chromogen. Overall, DSB had a more significant loss of miRNA.

### Shortened High Temperature Antigen Retrieval Improved the miRNA Yield

We noted differences in miRNA abundance at the HTAR step in our initial evaluation. Therefore, we varied the antigen retrieval time to 1 min, 10 min and 15 min, in an attempt to reduce the loss of RNA due to high temperature degradation. Each of these HTAR methods had an additional 22 min of heat up/cool down time, thus they represented 28, 32, and 42min of full experimental time. The intensity of staining for AE1/AE3 was equivocal at all HTAR lengths. The shortest HTAR had the most retained miRNA, with, on average 2.1-fold and 5.9-fold more at times 1 and 10 min respectively (Fig. 1b). Again, the different miRNAs had different fold increases. A second tested antibody, CD31, had insufficient staining with a HTAR of 1 and was evaluated by comparing 10 and 15 min of HTAR (Fig. 1c). The 10 min antigen retrieval had 6.9-fold more miRNAs compared to the 15 min using Let-7a, miR-222 and miR-128. Altogether, this data indicates that the shorter a feasible HTAR step can be, the higher the miRNA yield.

### RNAse activity was highest among primary antibodies and partially mitigated by RNAsecure

Next, the RNase activity of the IHC reagents was tested using an RNase alert system and measuring arbitrary units (AU) of fluorescence intensity in a CLARIOstar microplate reader. The negative control had a value of 340 AU and Millipore ddH20 water had a value of 325 AU. The highest value was for the AE1/AE3 antibody 5,330 AU. Three additional antibodies (Podocin, Claudin 8, CD34) had values between 680-966 AU. This indicated a need to reduce RNAse activity in the primary antibody incubation step.

In order to address the RNAse activity, we tested the addition of either 0.1% DEPC or RNAsecure to AE1/AE3+ IHC material and compared this with a spike-in of RNAse A. Both RNAsecure and 0.1% DEPC reduced RNAse activity (Fig. 1d). Additionally, the RNAsecure-treated material had the highest concentration of RNA (65 ng/μl) followed by 0.1% DEPC (20 ng/μl) and the untreated slides (15 ng/μl) remaining after AE1/AE3 staining.

We then evaluated the levels of three miRNAs from slides that underwent the full IHC protocol with or without RNAsecure relative to control unbaked slides. On AE1/AE3 stained slides, RNAsecure-treated experiments had an average of 24.6-fold miRNA loss relative to unbaked slides. The reduction in miRNA loss was variable across the three miRNAs (let-7a 8.8-fold, miR-128 4.9-fold and miR-101, 60.2-fold) (Fig 1e). The unprotected slides had a higher average fold loss (41.4) across the three miRNAs. The three measured miRNAs again had different fold changes (Let-7a 25.2-fold, miR-128 10.9 and miR-101 88.1). Thus, the use of an RNAsecure reagent had a mild, measurable improvement in mitigating RNA loss.

### RNAse activity in the chromogen also contributed to the loss of miRNA

The initial xMD-miRNA-seq was performed with a DSB chromagen to provide a black stain versus a brown stain of DAB. It was reasoned the darker color would improve pigmented cell transfer. However, DSB contains nickel and we became concerned this could have a deleterious effect on the RNA, due to the known effects of metal cations on DNA^17^. Therefore we compared DSB and DAB effects on RNA integrity. The fold changes in miRNA abundance were compared to slides not treated with chromogen. The DAB chromogen averaged a 4.7-fold loss, compared to a 10.6-fold loss for the DSB in comparison to an unstained slide (Fig. 1f). The fold change loss was variable across the three miRNAs for both DAB and DSB. Overall, DAB proved superior to DSB.

### Non-cross linked EVA membranes improved RNA yield

After the microdissection the cellular products must be extracted from the membrane, which is performed by dissolving the membrane in phenol:chloroform. The commercial 3M membrane is cross-linked, rendering it insoluble and yielding significantly lowering amounts of RNA. In order to improve miRNA yields after microdissection, a membrane using Elvax 410, a form of EVA, was generated and evaluated. We recognized these Elvax 410 membranes were “stickier” than the crosslinked membranes and impregnated fullerenes were used as a contaminant to break up the EVA polymer to provide greater specificity to the xMD method. To demonstrate this, heart tissue was stained with NCAD to highlight interacted disks (Fig. 2a) and xMD was performed. Fullerene impregnated EVA allowed for more specific capture of intercalated disks (Fig. 2b,c). Several different thicknesses of Elvax 410 membranes were evaluated for specificity and sensitivity (Fig. 2d-f). A thinner membrane captures less pigmented material while a thicker membrane captures more non-specific tissue. The 8% EVA membrane resulted in the best tradeoff of sensitivity and specificity in capturing pigmented cells and was the most consistent membrane to generate. The membrane thickness was determined by the gravimetric method (average 6.9 μm) and the film method (average of 5.21 μm) (Table 1).

**Table 1.**
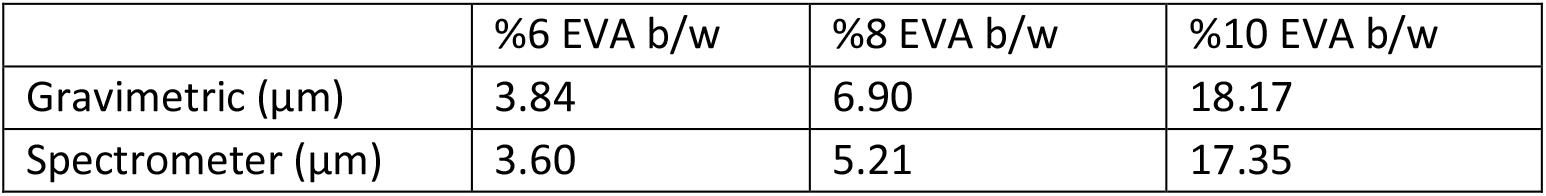
The thicknesses of membranes were tested gravimetrically and using a spectrometer.

**Figure 2.**
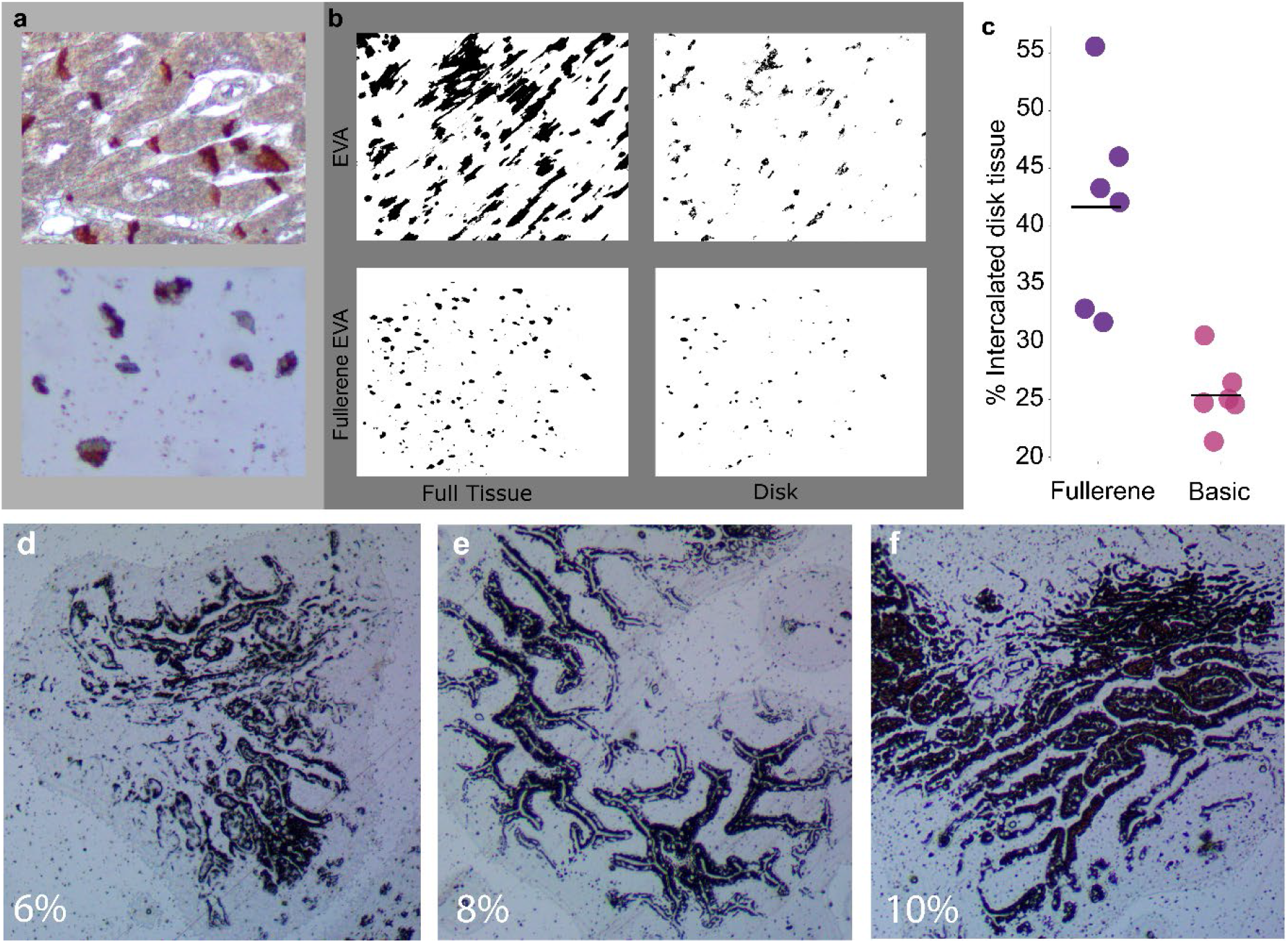
A comparison between fullerene-impregnated EVA membranes and unimpregnated membranes.**a**) Human heart slides stained for intercalated disks (NCAD) and cut at 4 μm were used. Top panel demonstrates the staining pattern for intercalated disks and bottom panel shows an EVA membrane. (**b**) Binary color area images of a post-dissected unimpregnated EVA membrane showing the full tissue captured (top left) vs the disk region within the collected tissue (top right) Similar images of a post-dissected fullerene-impregnated EVA membrane (bottom left) demonstrating improved capture specificity with less full tissue relative to captured disk material (bottom right). (**c**) An ImageJ pixel count across multiple images (N=12) demonstrated increase specificity of capturing intercalated disk tissue with fullerene-impregnated EVA membranes. (**d-f**) 6%, 8%, and 10% EVA membranes showing capture of AE1/AE3+ epithelial cells, with the cleanest pattern seen with 8% EVA.

RNA yield was determined for this optimal EVA membrane. Nucleic acid material was captured and assayed from slides stained with AE1/AE3 using 3M membranes, Elvax 410 membranes, or Elvax 410 membranes impregnated with fullerenes. The 3M membrane yielded the least amount of RNA of the three membrane types (9.4 ng/μl on average). The Elvax 410 produced the most (27.2 ng/μl on average) with the Elvax fullerene combination averaging 23.5 ng/μl, but with higher specificity.

### Methods to optimize specific RNA transfer to EVA membranes

Despite the optimizations, an initial small RNA sequencing experiment of xMD obtained AE1/AE3 positive cells failed to demonstrate appropriate gain or loss of epithelial (miR-192) and mesenchymal-specific (miR-143) miRNAs, suggesting unnoticed non-specific capture of RNA from the FFPE slide. We then performed experiments to improve RNA transfer specificity. Removing superficial RNA from the slide tissue by pressing an extra EVA membrane over the tissue prior to the flash lamp step improved the miR-143/miR-192 Ct ratio by 4.93. Lowering the energy (kJ) in the flash lamp step improved the ratio by 3.51. Combining these two steps further reduced the Ct ratio to 9.61 suggesting a 781-fold improvement in enrichment by the combined method.

### Tissue transfer is optimal with a glossy white background and high energy intensity

The flash lamp produces a quick, high intensity light which photothermally heats the pigmented areas to help affix them to the EVA membrane. Using the method described in the method sections, black, mirror, no background, white, and reflective white were tested. Both the black background (N=1) and no background (raised platform, N=1) qualitatively had poor transfer to the EVA membrane. The mirror background averaged roughly 1 million pigmented pixels (N=3), the matte white averaged 1.4 million pigmented pixels (N=5) and the glossy white background had an average of 1.7 million pigmented pixels (N=3) (Fig. 3a). We concluded a glossy white background was superior and would be used for all future experiments.

**Figure 3.**
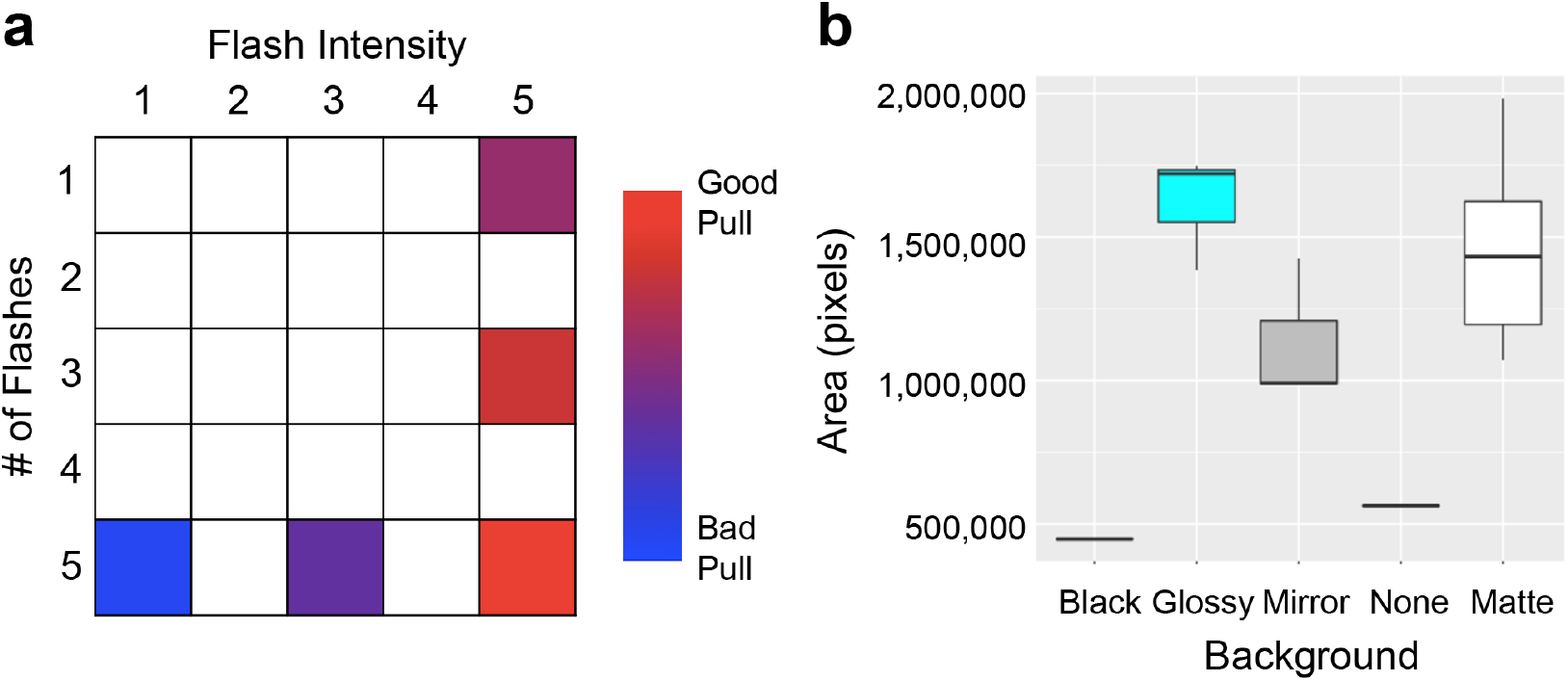
(**a**) Qualitative assessment of tissue yield from varying flash number and intensity. The higher intensity and most flashes resulting in the most material transfer. (**b**) Multiple backgrounds were tested during xMD. Quantitative comparison of tissue yield on membranes with a black, glossy white, mirror, none, and shiny white background. A glossy white background had the highest yield.

We then tested a select number of options of the flash lamp for intensity and number of flashes used, again evaluating AE1/AE3 staining in the small intestine. The most tissue transfer occurred under the highest intensity (level 5) and was independent of the number of flashes (1, 3 or 5). Less efficient transfers of pigmented material occurred at the lowest intensity (level 1) and the medium intensity (level 3) (Fig. 3b). All comparisons were qualitative in nature due to the clear differences observed.

### The fully optimized method has higher yields of miRNA before and after xMD

After all optimizations across the entire protocol, two comparisons were made. One compared the original and optimized protocols after all steps of the IHC method, but prior to the flash lamp capture of material. The second compared all steps through the full xMD protocol between the original and optimized protocols from material on the ELVAX 410 EVA membrane. For the pre-flash lamp experiment (scraped slide) on AE1/AE3 stained slides, the optimized method yielded an average of 39.9-fold more miRNA than the non-optimized method (Fig. 4a). The individual miRNAs again showed a difference in miRNA increase (Let-7a. 100.5 fold; miR-222, 6.2-fold; and miR-128, 11.4-fold). For the ELVAX 410 EVA membrane comparison of AE1/AE3+ material, there was, on average, a 16.45-fold greater yield in the optimized method relative to the original method, with some variation by miRNA tested (let-7a, 25.8-fold; miR-222, 13.7-fold; and miR-128, 9.9-fold) (Fig. 4b). The same experiments were performed with a CD31+ IHC to capture endothelial cells (Fig. 4c, d). Here the increases were individually 8.4, 14.4, and 29-fold improved for let-7a, miR-222 and miR-128 respectively in scraped tissue and 15-, 21.3- and 25-fold improved for let-7a, miR-222 and miR-128 respectively in xMD captured material.

**Figure 4.**
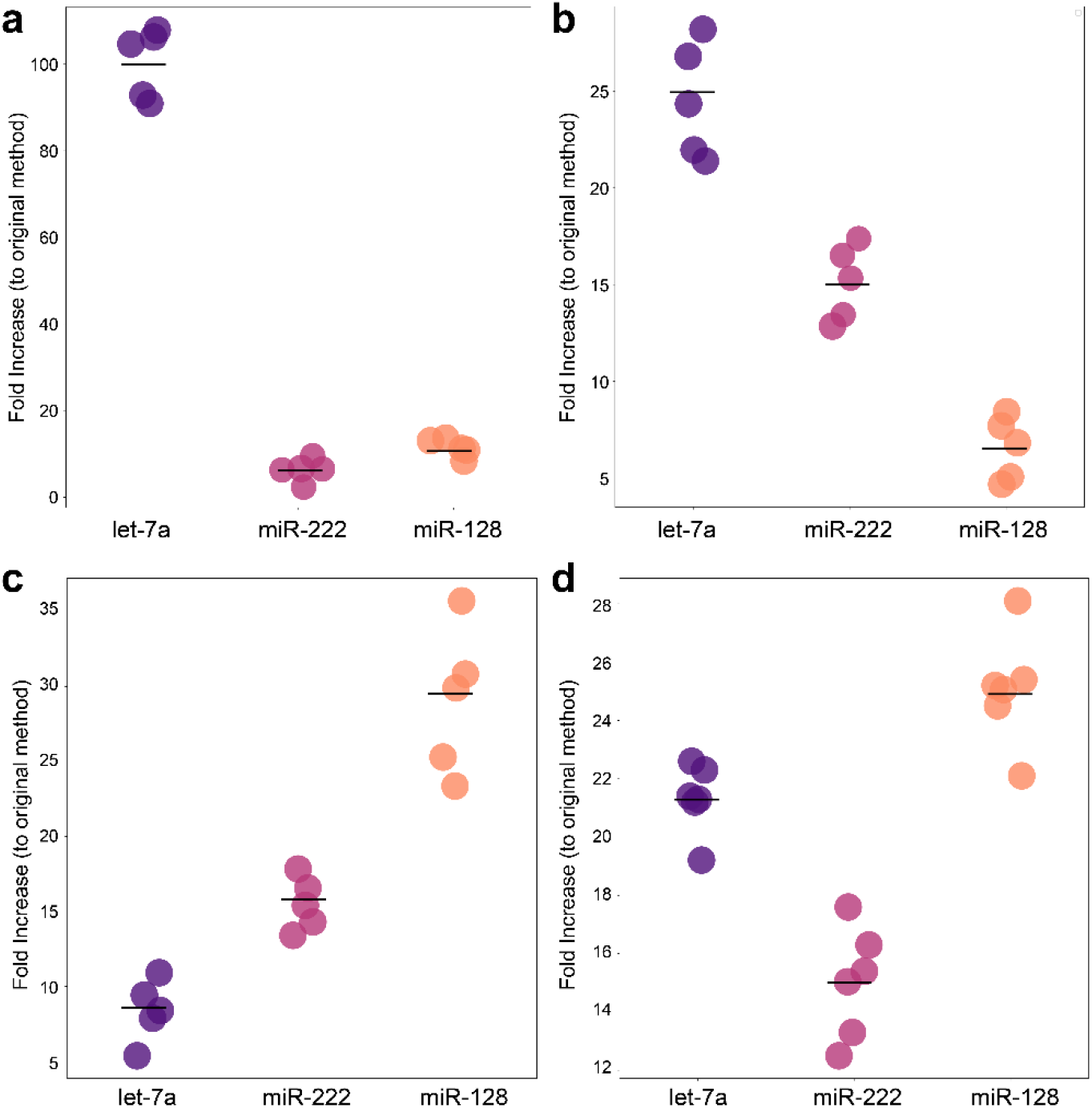
A comparison of miRNA yield from optimized IHC and xMD to original methods.(**a**) miRNA yield directly from slides which underwent AE1/AE3 IHC. Figure shows the fold improvement of the optimized protocol to the original protocol (N=5). (**b**) miRNA yield after xMD capture of AE1/AE3+ material. Figure shows the fold improvement of the optimized protocol to the original protocol (N=5). (**c**) miRNA yield directly from slides which underwent CD31 IHC. Figure shows the fold improvement of the optimized protocol to the original protocol (N=5). (**d**) miRNA yield after xMD capture of CD31+ material. Graph shows the fold improvement of the optimized protocol to the original protocol (N=6).

### A TLDA qPCR array demonstrates enrichment of AE1/AE3+ cells

After all optimizations, TLDA miRNA array was performed on duodenal tissue scraped from the slide or from AE1/AE3 positive xMD material, to document global miRNA changes in the epithelial cell specific population. The TLDA miRNA array contains 370 miRNAs, of which 258 miRNAs had expression that could be analyzed. TLDA was chosen as it could be combined with a QC step directly on the same RNA sample, due to the low input RNA requirement. Thus, prior to the TLDA, QC was performed to compare the ratio of miR-192 to miR-143 between AE1/AE3+ cells and a tissue scrape using standard qPCR. This method demonstrated a 29.9-fold relative increase in miR-192 to miR-143 between AE1/AE3+ epithelial cells and the total tissue scrape, indicating enrichment was made and the TLDA proceeded. The TLDA qPCR Ct data was normalized, and fold changes between the whole tissue and epithelial cells were calculated. Epithelial cells represented ∼15% of the cellular material in the duodenal slide, which indicates potential maximal gains and losses in miRNA relative to the full tissue sample. Globally, 54 miRNAs had >4-fold decreased expression in epithelial cells, while only 14 were >4 fold enriched. Two main mesenchymal markers, miR-143 and miR-145, were 336- and 279-fold reduced in the epithelial cell sample (Fig. 5a). Conversely, the miR-200 family of epithelial markers, increased between 1.7-fold and 14-fold in the epithelial cell material (Fig. 5b). Two additional miRNAs increased in the epithelial cells were miR-488 and miR-302c (9.6- and 7.1-fold respectively).

**Figure 5.**
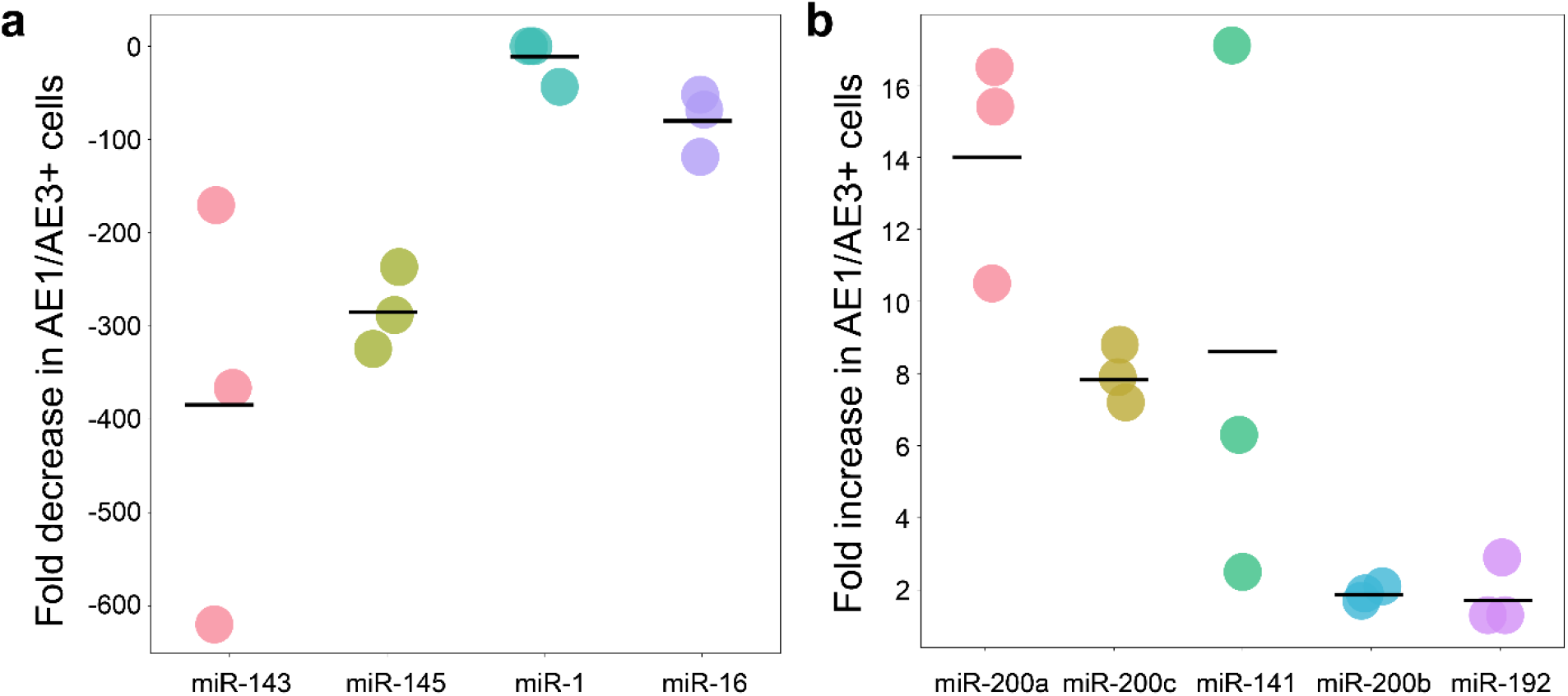
TLDA array results (**a**) Selected miRNAs that had notably lower expression in AE1/AE3+ epithelial cells relative to the total tissue. (**b**) Selected miRNAs that had notably higher expression in AE1/AE3+ cells relative to total tissue.

## Discussion

The original publication of the xMD-miRNA-seq method demonstrated the potential utility of a method to capture near *in vivo* miRNA expression patterns of specific cells^14^. However, the method was inefficient with notable RNA loss during the xMD steps and a low yield of miRNAs in the sequencing step. Thus, it was impractical for widespread usage. Here we assessed each step of the xMD-miRNA-seq method to reduce RNA loss and increase the specificity of miRNAs transferred onto the EVA membrane. The key improvements were realized by adding an RNAse inhibitor, shortening the HTAR time, using a nickel free chromogen, generating a non-crosslinked fullerene-impregnated EVA membrane, and using a phenol:chloroform extraction method. These changes increased RNA yield, of all three examined miRNAs, individually as well as combined. The qPCR fold changes showed an increase in miRNA collection, averaging 16.5-fold when all changes were combined. Through these improvements we were able to obtain ∼200 ng of total RNA per slide, and ∼1 μg when five slides are combined. As newer small RNA sequencing methods can be performed with as little as 10 ng of starting RNA, these improvements greatly expand the opportunity of using this method across rarer and more specific cell populations.

These optimizations, however, did not result in improved specificity of the collected RNA. Thus, a separate set of optimizations resulted in a marked reduction of ambient RNA from the surface of the slides and reduced non-target transfer of material. These steps were needed due to the stronger transfer properties of the non-crosslinked EVA membranes compared to previously used crosslinked EVA membranes.

A limitation of the method was our inability to execute the full assay with small RNA sequencing. Two attempts ably provided a robust miRNA dataset (>580 bona fide miRNAs), but neither had the appropriate enrichment expected^18,19^ The first was performed before the corrections to the protocol (E – xMD specificity optimization) and had no meaningful enrichment. The second was performed after developing a QC step, but the QC was not done concurrently. This sample had a ∼50% enrichment in epithelial cells based on gains/losses of expression of epithelial and mesenchymal markers. Neither assay was performed with a concurrent qPCR QC step due to the relatively high need of RNA for both routine qPCR and sequencing library preparation protocols. Utilizing a QC step prior to performing the global miRNA array ensured the quality of the xMD, which we now recognize as critical prior to genomic level assaying and must be included in future iterations of this method that employ sequencing. Nonetheless, using all of the optimized approaches, we noted a strong enrichment for epithelial cells by the qPCR array-based method.

We demonstrated the utility of the xMD method with a 4-13 fold increase in epithelial-specific markers (miR-200 family), and more importantly, a >300 fold decrease in mesenchymal miRNA (miR-143/145) expression compared to the entire duodenal tissue^4,20^. Of note, AE1/AE3 is a pan-cytokeratin marker of all epithelial cells of duodenum from the stem cells at the crypt base, through progenitor cells, to the mature enterocytes at the top of the crypt. This may explain why a couple of miRNAs known for their specificity to stem cells (miR-488, miR-302c) were also enriched^15^. Future uses of xMD will employ IHC antibodies selected for more specific labeling sub-groups of the epithelium based on single cell and proteomic expression data^8,21-24^.

Not only was the IHC and miRNA extraction optimized, the EVA membranes used for xMD were optimized. We developed non-polymerized EVA membranes, with lattice disruption, which were able to partially dissolve in phenol:chloroform allowing for easier extraction of cellular products and higher yield.

While there is much excitement for single cell RNA sequencing methods to assay miRNAs, challenges remain in that domain as well. First, miRNAs represent a very small fraction of total RNA in a given cell, thus the yield per cell is low and obtaining that limited information on a cell-by-cell basis is cost-ineffective. Secondly, at the low yield per cell, the ability to identify the full repertoire of miRNAs (low to modestly expressed) in a cell type is challenging to impossible by that approach^25^. Finally, the great strength of single cell RNA sequencing is to define populations of cells, however, based on our experiences with the limited set of miRNAs that exist (relative to genes), miRNAs will be inferior to genes for this purpose.

The optimized xMD method, described here, can allow any researcher access to *in vivo* miRNA expression patterns from cells of interest from FFPE tissues. Between single cell human cell atlas projects and the human protein atlas, a large number of cell-type defining genes/proteins are being realized that can further refine the selectivity of this xMD method (cite single cell papers^8,21,22,26,27^. Additionally, most materials used in this optimized version of xMD are widely available and cost-effective. Those that are not, require little training to make. Other methods of microdissection, such as laser capture, are limited by the ability of users to obtain enough specific material and/or access to expensive machinery. The smaller expense and experience required make xMD ideal for a new method of microdissection.

This new optimized method will allow for more near *in vivo* miRNA expression patterns of cells to be determined. Previous work has shown miRNA expression *in vitro* differs from *in vivo* cells, highlighting the importance of the *in vivo* environment in FFPE tissues. miRNAs are effective in the cells in which they are expressed and this population of miRNAs varies from cell type to cell type. The expression profiles of individual cells are lost when whole tissue expression is performed, limiting our current understanding of miRNA expression in cell types.

In conclusion, we demonstrate a fully optimized xMD method that obtains significant amounts of highly specific miRNA from cells obtained from FFPE tissues. This method showed robust isolation of epithelial cells from duodenal tissue, establishing an *in vivo* expression pattern of this cell type.

## Acknowledgements

This work was supported by the National Institute of General Medical Sciences, R01GM130564 and the National Heart, Lung, and Blood Institute, R01HL137811 to M Halushka. We thank the James Segars laboratory for the use of the ClarioSTAR fluorometer, the Ben Larman lab for the use of the qPCR thermocycler, the Oncology Tissue and Imaging Services Core for slide preparation, Margaret Weinstein and Mohammad Dbouk for sample acquisition, Arun Patil for figure assistance, Anandita Rajpurohit and Joo Heon Shin for library preparation assistance, and Roxane Ashworth and the GRCF DNA services staff for TLDA plate discussion and assistance.

## Author Contributions

AEJ performed the main experiments and wrote the manuscript; BB and KAPM performed optimization experiments; CO, DC and JAB assisted with sequencing; KMJ assistance with data interpretation; DM and DHK assisted with membrane analysis, MHR oversaw membrane development; AZR conceived of the project, helped oversee the project and assisted on the manuscript; and MKH oversaw the project and helped write the manuscript.

## Data availability

TLDA array data is available at the Gene Expression Omnibus, study GSE205216.

